# Landing manoeuvres predict roost-site preferences in bats

**DOI:** 10.1101/2022.03.12.484101

**Authors:** Gloriana Chaverri, Marcelo Araya-Salas, Jose Pablo Barrantes, Tere Uribe-Etxebarria, Marcela Peña-Acuña, Angie Liz Varela, Joxerra Aihartza

## Abstract

Roosts are vital for the survival of many species, and how individuals choose one site over another is affected by various ecological factors. Biomechanical constraints could also affect roost selection, particularly in volant taxa that require sites with easy access, thereby reducing costs (i.e., predation, accidents). To date, no studies have established an association between landing performance and roost-site selection, as predicted by biomechanical constraints associated with flight. We aim to determine roost-site selection in disc-winged bats (*Thyroptera tricolor*), a species known to roost within developing tubular leaves. This study is coupled with various experiments that measure how a conspicuous apex affects landing tactics and performance. We show that *T. tricolor* prefers leaves with a longer apex, the space typically used for landing. Bats also approach and enter these leaves more consistently, increasing task performance while reducing the risk of injuries.

**Summary statement:** Spix’s disc-winged bats prefer to roost in some types of leaves, which we show may be related to costly maneuvers during the approach and landing phases.

## INTRODUCTION

The relationship between living beings and the resources they require for survival and reproduction represents one of the core topics in ecology. The most critical resources needed by animals to secure their survival include food and refuge, and our understanding of how and why species select among several options has been strongly rooted in optimality models that predict animal’s choices based on cost-benefit analyses (Pyke, 1984; Rapport, 1971). However, we know that the decision to select one resource over another is also strongly influenced by various constraints imposed by a species ecological context and its behaviour and morphology (Brost et al., 2015; Machado, 2020; Shakeri et al., 2021). For example, a particular animal will consume a given food item if it’s found within its range, competition for it is relatively low, resource production or availability matches its daily activity periods, and the animal has appropriate morphological structures that allow it to obtain and process this food item efficiently. Knowing the interactions among these constraints allows us to understand why major resource categories are selected (e.g., nectar vs fruits) and how animals select species within those categories, or individuals within species. The latter is of particular interest because it provides the basis for natural selection in both antagonistic and mutualistic interactions (Andreazzi et al., 2017; Law, 1985; Nuismer et al., 2013; Thompson, 1988).

While many studies to date that tackle resource use have focused on the relationship between animals and their food sources, a key component of many species’ natural history are roosts. Roosts provide sites for animals to escape predation and inclement weather conditions and to conduct crucial fitness-related activities such as copulation and lactation (Kunz, 1982). The selection of roosts is known to be influenced by various factors, including temperature, humidity, access to feeding sites, or increased protection against predators (Barros et al., 2020; Boonman, 2000; Delancey and Islam, 2020; Kerth et al., 2001; Rogers et al., 2006). However, other aspects of a species’ biology can limit the range of roosting resources that can be exploited, including biomechanical constraints. In bats, biomechanical constraints are known to limit the use of different categories of roosting resources (Boerma et al., 2019; Riskin et al., 2009), but they may also influence the selection of specific roost-sites within categories, as biomechanics are tightly linked to manoeuvrability in volant taxa (Liu et al., 2016; Roderick et al., 2017; Tobalske, 2007), which in turn may determine access to some resources but not others. Studies that have gauged tree-cavity selection in birds and bats, for example, often use accessibility arguments to explain roost-site preferences; essentially, cavities that provide easy access reduce flight costs and the risk of predation (Fisher et al., 2004; Vonhof and Barclay, 1996). Given the energetic cost of flying (Norberg, 1996), the risk of accidents during landing events (Wheatley et al., 2021), and that the process of roost switching can render many animals vulnerable to predation (Speakman et al., 1994), rapid and safe access may be a particularly relevant factor that explains roost site selection. Notwithstanding, no studies to date have quantified this association between landing performance and roost site selection, as predicted by biomechanical constraints associated with flight.

An interesting case of an extreme specialization for a specific roost type occurs in Spix’s disc-winged bats, *Thyroptera tricolor*. This species shelters inside the developing tubular leaves of plants in the order Zingiberales (Findley and Wilson, 1974; Vonhof and Fenton, 2004). To attach themselves to the smooth inner surfaces of the tubular structure, they have discs on their hands and ankles that provide adhesion primarily through suction (Riskin and Fenton, 2001; Schliema, 1970; Wimsatt and Villar, 1966). This bat is strongly dependent on this type of roost because their reduced thumbs preclude them from effectively clinging to rough surfaces as most bat species (Riskin and Fenton, 2001), and experimental removal of potential roost sites shows that they seem incapable of using alternative roosting structures (Chaverri and Kunz, 2011). A recent study by (Boerma et al., 2019) also shows strong biomechanical specializations for using these furled leaf roosts. Specifically, they show that the force with which bats land onto the leaves is significant and that the discs are vital for effective attachment to the leaf. Their results also suggest, albeit this was not directly tested nor discussed, that leaf shape can affect the value of different plant species for roosting since bats appear highly reliant on the leaf’s long apex, a structure that is not present in all plant species used by this bat, for safe landing and rapid post-landing settlement.

This study aims to determine if biomechanical constraints influence roost-site selection in *Thyroptera tricolor*. Previous studies suggest that this bat prefers certain plant species and leaf shapes for roosting (Chaverri and Kunz, 2011) and that these preferences may relate to differences in microclimatic conditions within the leaves and reduction of conspicuousness as a mechanism to reduce predation pressure (Pérez-Cárdenas et al., 2019; Solano-Quesada and Sandoval, 2010). However, given the results of Boerma et al. (2019b), we expect that biomechanical constraints could also be highly relevant during the process of roost-site selection. Our study provides a thorough understanding of leaf-shape preference in natural populations coupled with various experiments that measure how the presence of a conspicuous apex affects landing tactics and performance. Overall, we predict that i) bats will show a significant preference for plants with an acuminate apex over truncated leaves (short apex); and that ii) landing will be consistently more effective (according to the landing patterns recorded by Boerma et al. (2019b) in leaves with an acuminate apex. Our experimental results provide a strong argument for biomechanical constraints playing an essential role during roost site selection in bats.

## MATERIALS AND METHODS

### Plant species preferences

We performed systematic surveys of furled leaves at 6 study sites (Bolivar, Esquinas, Finca, Lecheria, Naranjal, Ureña) in lowland tropical forests in southwestern Costa Rica. We searched for leaves that could be used as potential roost sites by *T. tricolor* an average of twice a month from October 2006 through July 2007. Surveys increased in frequency (up to once every week) during the parturition period (February – April 2007), and some sites (i.e., Naranjal) were surveyed more often due to low recapture rates. During the first 15 surveys, we quantified the number of furled leaves (i.e., potential roosts) per site by counting the number of unoccupied and occupied (i.e., roosts) leaves. Unoccupied furled leaves were only counted if their opening diameter ranged 4 to 20 cm, following Vonhof & Fenton’s (2004) results. However, occupied leaves were counted even if their opening diameter was not within that range.

We used 104 sampling events from our 6 study sites. The number of available potential roosting leaves was compared against the number of roosts (used leaves) for the two most commonly used plant species at our study sites: *Heliconia imbricata* and *Calathea lutea*. Bayesian generalized linear models were used to evaluate which parameters explain roosting leaf occupancy better (proportion of available furled leaves used as roosting sites). The number of used roosts and total available tubular leaves by plant species was used as a combined response variable modelled with a binomial distribution. We evaluated three models: one includes species and density as predictors, another one only includes species, and the last only includes density. All models include the site as a random effect.

### Leaf-shape preferences

To characterize leaf shape, we measured leaf height, leaf length, tip length, and tip and mid circumferences for undamaged roosts after capturing bats (Figure 1a). Based on these measures, we also calculated available space (leaf length minus tip length), the ratio of tip length to tip circumference (tip length divided by tip circumference), and roost height (leaf length plus leaf height). However, available space and roost height were excluded from further analyses due to high collinearity. Supervised Random Forest (Breiman, 2001) was used for discriminating species based on leaf shape parameters (detailed below). A randomization test on variable importance was used to identify those parameters significantly contributing to model discrimination.

**Figure 1.**
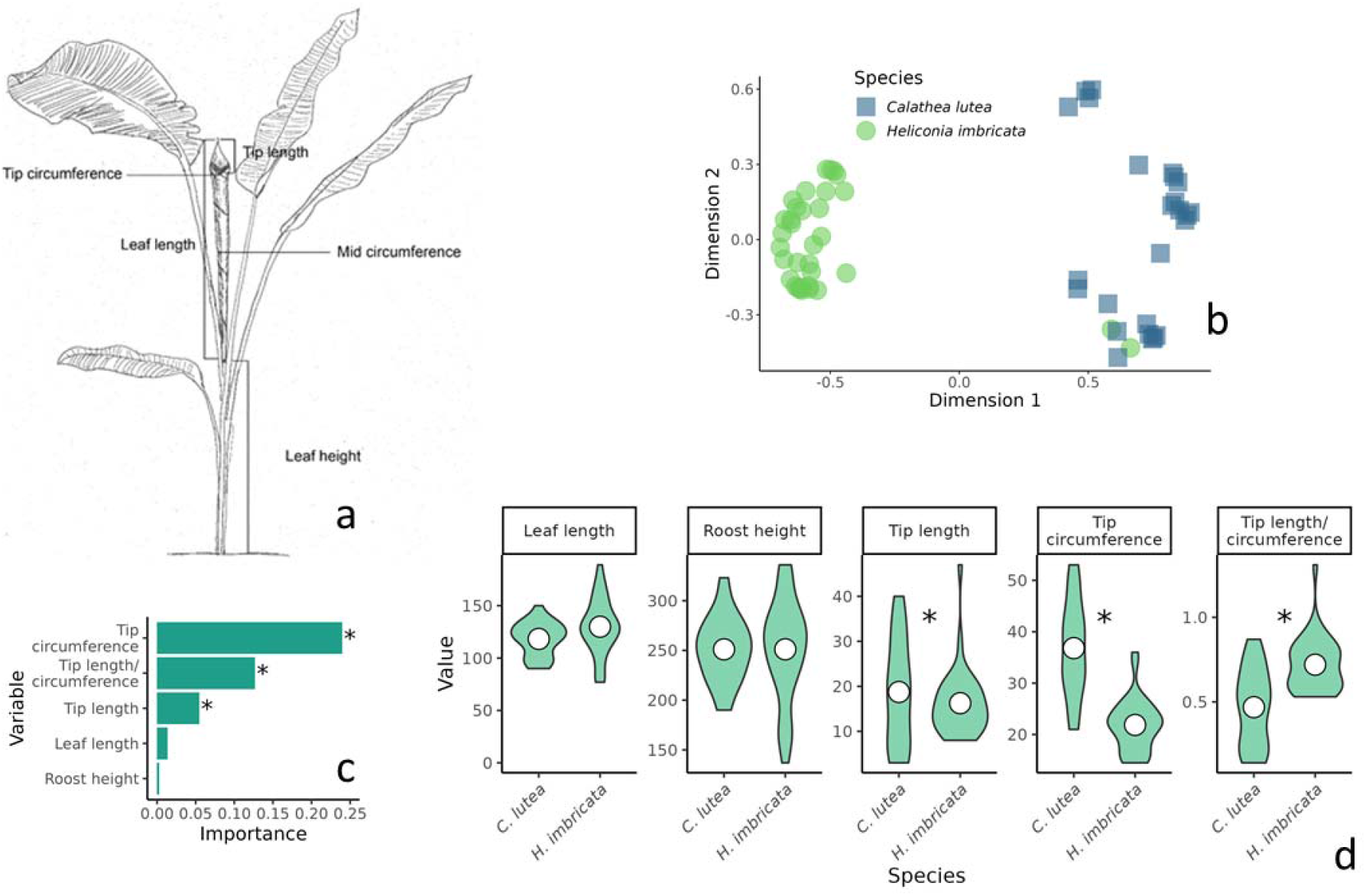
(a) Morphological parameters recorded from roosts. The tip circumference was measured at the tube opening, and the mid circumference was measured at the centre of the furled leaf (i.e., leaf length/2). In addition to these measures, we also calculated roost height — the sum of leaf height and leaf length— and available space —the difference between leaf length and tip length—. (b) Leaf shape space based on Random Forest proximity. (c) Random Forest importance (Gini impurity units) of the five variables used to classify furled leaves of *H. imbricata* and *C. lutea.* Higher values imply a higher contribution for discriminating between the two species. (d) Violin plots compare morphological variables between *H. imbricata* and *C. lutea*, asterisks denoting a significant difference in a given parameter between species. Values for all traits, except tip length/circumference, are in cm.

### Flight manoeuvres while entering leaves

We experimentally evaluated variation within flight manoeuvres employed for entering furled leaves of variable tips or apex shapes. Two types of apex shapes were used, representing the two extremes observed in the leaves used as roosts by *T. tricolor*: a pointy tip (i.e., acuminate apex) extended above the opening of the leaf (~ 5 cm tip length) and a leaf completely truncated above the opening of the leaf (i.e., no extended leaf area after the opening; 0 cm tip length). Single individuals were released inside a flight cage containing a plastic furled leaf of one of the two types (acuminate or truncated). Flights were videotaped using two digital cameras at 60 fps (SONY HDR-XR160). Flight trajectories in tri-dimensional space were then inferred from videos using PhysMo Video Motion Analysis v2.0, Microsoft Excel and SketchUp (2016).

We characterized flight manoeuvres on two time periods prior to entering the leaf: last inflection period (a variable period after the last inflection on the z-axis) and ballistic descent period (a fixed period of 11 ms before landing, as described by Boerma et al. (2019b). We measured four parameters at the start of each period: height, acceleration, distance to the leaf’s opening and vertical angle. We also included the number of height inflections during the ballistic descent period. Differences in leaf landing manoeuvres between acuminate and truncated leaves were evaluated using supervised Random Forest (Breiman, 2001) on the described parameters. We used a Monte Carlo randomization approach (e.g., randomization that does not necessarily generate all possible combinations; Robert & Casella 2010) to test the statistical significance of the discrimination. To do this, we built a routine on shuffling the leaf-type categories to unlink them from any structure in the parameter data and calculated the out-of-bag (classification) error, which was replicated 10 000 times. Discrimination errors obtained from the randomization procedure were then compared to the observed value on the original data set. We calculated the p-value as the proportion of expected random values lower than the observed value (i.e., how likely it was to obtain the observed discrimination by chance). In addition, we evaluated differences in multivariate variance in flight manoeuvres for the two apex shapes using an analysis of multivariate homogeneity of group dispersions (Anderson, 2006) in the R package vegan (Dixon 2003).

### Landing performance

To determine if the presence of a conspicuous leaf tip facilitates landing, we registered the landing patterns of individuals onto acuminate and truncated tubular leaves. For this, we allowed bats to fly inside a flight cage for a maximum of 5 minutes or until they entered a leaf. The individual bats were tested on each leaf type once. The order in which the leaf types were presented to bats was randomized. Landings were recorded with a GoPro Hero7 Black camera (GoPro Inc., California). Videos of bats’ landings were analysed to determine 1) whether bats landed first with the thumb disks and then attached the foot disks (Boerma et al., 2019), and 2) the time elapsed since the bat made first contact with the leaf until it descended beyond the leaf’s rim. We believe the latter is relevant as it provides an estimate of how long it takes the bat to become inconspicuous to potential predators. For the former, landings that followed that pattern were considered “normal”, whereas those that did not were considered “odd”. A Bayesian generalized regression model was used to assess if landing pattern (normal or odd) was associated with apex type (acuminate or truncate). Bernoulli distribution and a logit link function were used to model the response variable. We also evaluated the link between apex type, landing pattern and the hiding latency inside the furled leaf. For this, we use Bayesian mixed model regressions with latency as the response variable, individual as a mixed factor, and either apex type of lading pattern as predictors. A single multiple regression model was not evaluated as both predictors were found to covary (see results), which precludes inferring their effect on the third variable in a single model.

### Statistical analyses specifications

Supervised Random Forest models (Breiman, 2001) were run using the R package ranger (Wright & Ziegler 2017). Optimal tuning parameters for Random Forest models were estimated in the R package tuneRanger (Probst et al. 2019), using 10000 trees and 1000 iterations. All parameters were Box-Cox transformed previous to analysis, and highly collinear variables were excluded (absolute r > 0.9). Out-of-bag error (the mean prediction error on each sample using only the random forest trees that did not have that sample) was used as a classification performance metric. Random forest importance was used to assess the relative importance of predictors, and importance was calculated using Gini impurity (Breiman, 2001).

All regression models were run in Stan (Stan Development Team, 2021) through the R platform (R Core Team, 2021) using the R package brms (Bürkner, 2017). All continuous predictors were z-transformed to remove differences in magnitude and simplify interpretability. We present effect sizes as median posterior estimates, and 95% credibility intervals (CI) as the highest posterior density interval. Parameters in which credible intervals did not include zero were regarded as affecting the response variable. We applied a model averaging approach for parameter estimation on plant species preference in which several models were evaluated. We used the Bayesian leave-one-out information criterion (LOOIC, Vehtari et al., 2017) with the R package loo (Vehtari et al., 2020) to assess the relative support of models to the data. LOOIC weights, which quantify the relative support compared to other candidate models, were then used to calculate weighted estimates averaged across models (i.e., model averaging).

All models were run on three chains for 30 000 iterations, following a burn-in of 3000 iterations. The adequate sample size was kept above 3000 for all parameters. Performance was checked visually by plotting the trace and distribution of posterior estimates for all chains. We also plotted the autocorrelation of successive sampled values to evaluate the independence of posterior samples. A potential scale reduction factor was used to assess model convergence and kept below 1.05 for all parameter estimates.

## RESULTS

During our systematic surveys, we recorded a total of 2,715 furled leaves comprising seven plant species: *Calathea inocephala*, *C. lutea*, *Heliconia imbricata*, *H. irrasa*, *H. latispatha*, *H. stilesii*, and *Musa* sp. The two most commonly used plants for roosting were *H. imbricata* and *C. lutea* (Figure 2a). These two species were also the most commonly recorded plants with furled leaves (Figure 2b). Furled leaves were most commonly observed in *H. imbricata* at all sites except for Ureña, where the most common species with furled leaves was *C. lutea*. Nonetheless, plant species, but not plant density, significantly predicted the usage of furled leaves as roosts by *T. tricolor* across sites (Figure 2c; species effect size: 1.30, CI: 0.94–1.66; density effect size: 0.0003, CI: −0.01–0.01).

**Figure 2.**
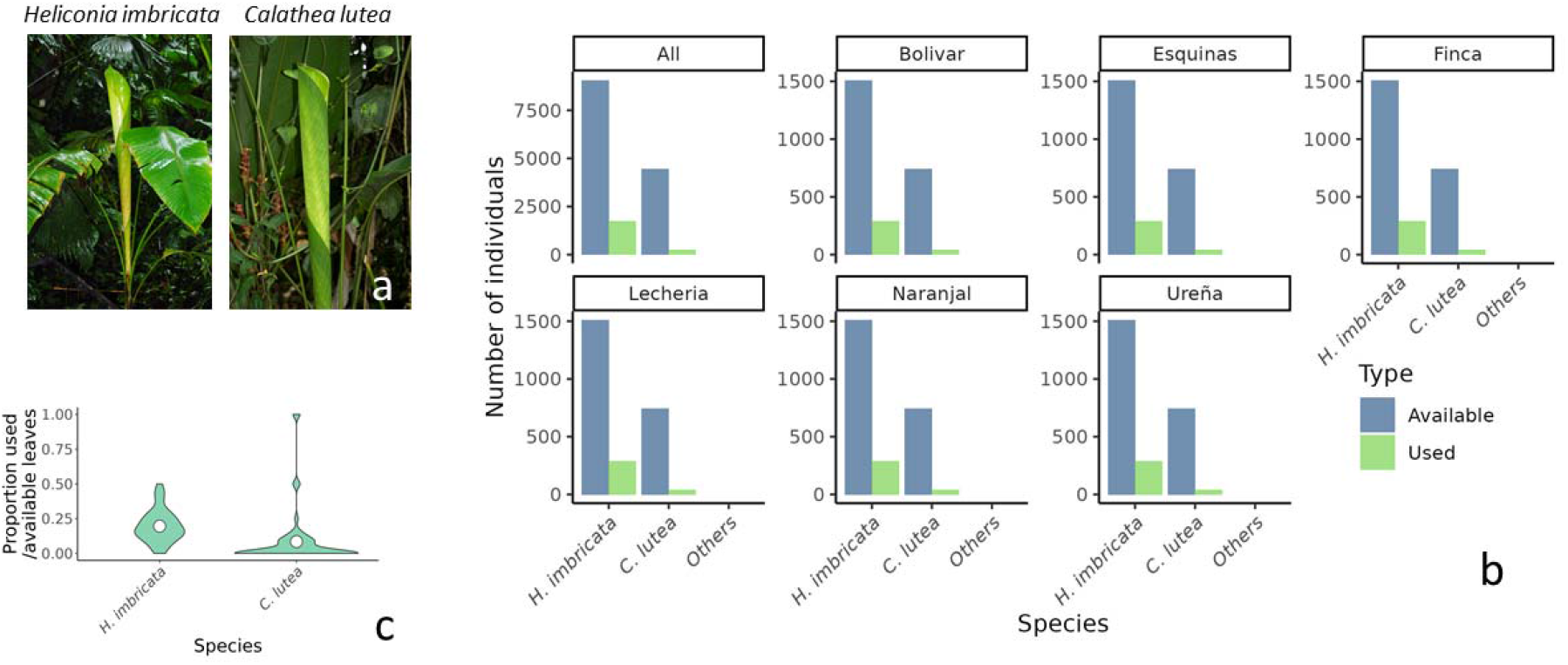
(a) Photographs of the two most commonly used plant species by *Thyroptera tricolor*, *Heliconia imbricata* and *Calathea lutea*. (b) The numbers of used and unused (available) furled leaves per plant species. Detailed results are presented only for the two most commonly used plant species, while less commonly used plants are grouped into a single category. Data are shown for each site separately and also collectively. (c) Violin plots of the proportion of used per available furled leaves.

A Random Forest model classified the furled leaves of *H. imbricata* and *C. lutea* with an out-of-bag error of 3.38% (i.e., 96.6% correctly classified), indicating leaf shape differences between the two species (Figure 1b). Only leaf tip parameters (tip circumference, tip length to tip circumference ratio, and tip length), but not overall leaf parameters (leaf length and roost height), contributed significantly to species discrimination (Figures 1c and 1d).

Flight trajectories were classified according to the apex type of the leaves they were landing on for 73.7% of the trials (Random Forest out-of-bag error of 26.3%), and this classification was significantly higher than expected by chance (p = 0.006). Multivariate variance was significantly higher on flight trajectories from experiments using truncate leaves (F = 6.80, df = 1/36, p = 0.0132; Figure 3a). The landing pattern was associated with apex type. Odd landings were estimated to be more than three times more common in truncate than acuminate furled leaves (Figure 3b; effect size: 3.28, CI: 1.01–6.23). No effect of landing pattern (effect size: 205.51 CI: −88.54–495.01) or apex type (effect size: 31.52, CI: −245.13–306.49) was detected on latency to hide (Figure 3c).

**Figure 3.**
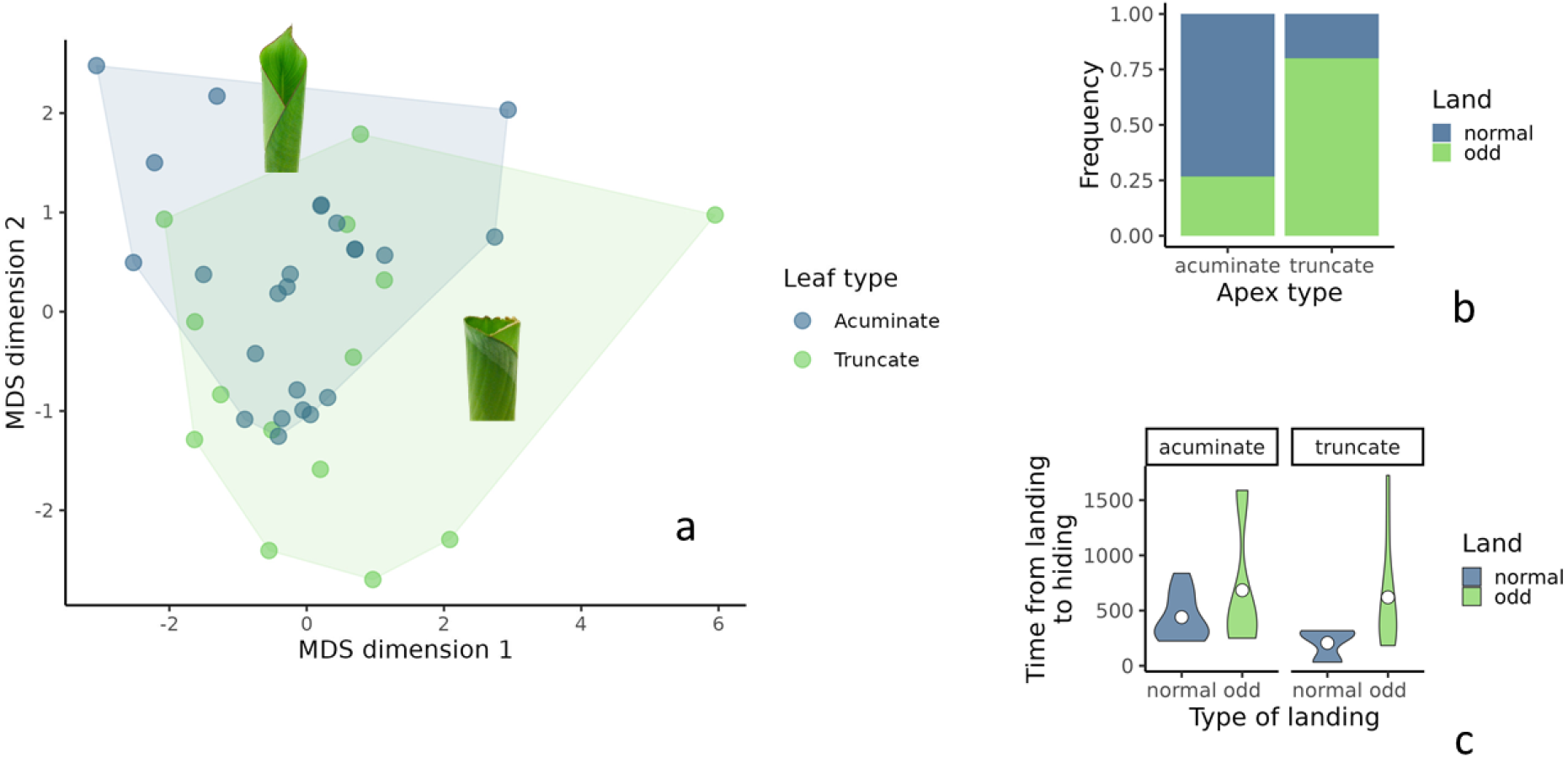
(a) Dispersion of estimated flight trajectories during approaches towards acuminate and truncated leaves (photos of each type included), projected using multidimensional scaling (MDS). (b) The proportion of regular and odd landings on the tubular leaf types (acuminate or truncate) used in the experiments. (c) Time elapsed since bats first made contact with the leaf’s structure until they moved beyond the rim.

## DISCUSSION

Our study shows that *T. tricolor* exhibited a significant preference for furled leaves in *H. imbricata* over *C. lutea*, which were the two most common plant species sampled at our study sites in southwestern Costa Rica. This preference is not related to overall availability but is most likely linked to morphological differences between the furled leaves produced by these two species. The most important traits that separate roosts selected in these two plants correspond to features related to the leaf space used for landing when bats reach the roost: length of the leaf’s apex (longer in *H. imbricata*), the leaf’s width (broader in *C. lutea*), and the relationship between these two characteristics.

The observed differences in morphology between the two plants most commonly used by *T. tricolor* may affect bats in two different ways. A narrow leaf probably allows bats to remain inconspicuous to diurnal predators that search for bats from above, such as monkeys and several species of birds of prbats’ preference for roosting in *H. imbricata*, as narrow leaves are available in both species. Another possibility is that bats select the plant species that provide a longer-lasting tubular structure, thus decreasing the energetic and predation costs involved in the location of a new roost during the daytime. Previous studies show that tubular leaves in some *Heliconia* species, compared to *Calathea*, last longer (up to 31 hours) within the bats’ preferred circumference (c.a., 22.30 cm; Vonhof & Fenton 2004; ey (Boinski and Timm, 1985), since less light will enter the tube and bats usually roost at the bottom. Considering that avoiding predation is of utmost relevance, it makes sense that tip circumference is the main characteristic preferred by bats when selecting roost-sites. However, this trait alone cannot explain the Pérez-Cárdenas et al. 2019a). Notwithstanding, tubular leaves in *Calathea* species might still be available within the bats’ preferred circumference for up to 14 hours (Vonhof and Fenton, 2004). Considering that *T. tricolor* changes roosts daily, this difference in roost longevity does not seem particularly significant in explaining the bats’ preference for roosting in *H. imbricata*, yet certainly seems to deserve further examination.

Another way plant morphology may influence bats is by determining how they should approach and land on tubular structures. *T. tricolor* has the challenging task of rapidly entering a tubular structure whose opening is scarcely broader than the bat itself. This task seems greatly facilitated by the presence of an apex, which bats typically use as the first contact with the roost, aided by the suction of their discs. Bats land on the roost in various ways without an apex, including the first contact between forearms (and other body parts) and the leaf’s rim. While these odd landings do not seem to increase the time needed for bats to hide within the tubular structure (potentially avoiding being detected by predators), they might cause injuries or, occasionally, failed entries. The impact force applied by *T. tricolor* upon landing on leaves is very significant (mean peak total impact force = 6.98 bodyweights), the strongest measured in bats until date (Boerma et al., 2019). This impact force could be enough to damage bones if the force is not dampened sufficiently or the direction of contact is not appropriate, so being able to land in a controlled fashion may be critical for a bat’s safety. In this respect, it is well known that bones are more resistant to fractures when forces are applied longitudinally, not transversely (Behiri and Bonfield, 1989; Li et al., 2013), as occurs for forearm bones when *T. tricolor* lands in a usual way. Thus, the probability of bone fractures could increase if bats’ forearms land first on the leaf’s rim since this would represent a transverse force. Also, being able to slow down upon entering the leaf, aided by the attachment provided by the suction discs onto the leaf’s inner wall, could be important if other group members are already roosting within it. An apex presence may also allow bats to quickly confirm that they can land on this surface without hitting group mates.

The absence of an apex may also influence a bat’s approach to the roost’s entrance. We found that flight manoeuvres of *T. tricolor* differ depending on the apex morphology of the roost leaf and that manoeuvres were more variable when approaching truncated apex leaves. In many motor tasks, especially when these are dangerous or require a long learning period, animals exhibit highly stereotypical motor patterns and movement trajectories. Performance in a motor task is often facilitated by reducing variability in the actions required to conduct that task (Dhawale et al., 2017). Given the potential costs of abnormal landings, a more defined landing strategy for approaching acuminate tubular leaves could result in a lower risk of injury when approaching such roosts, suggesting that a biomechanical constraint might play a role in roost choice in *T. tricolor*.

In conclusion, we postulate that both ecological and biomechanical constraints strongly influence roost-site selection in *T. tricolor*. Bats might select narrow leaves with a long apex to avoid predation and reduce the potential biomechanical costs involved in entering leaves with no safe landing surface. These findings add another layer of understanding of the species’ complex interactions with the resources it requires for survival. We know that the degree of resource specialization is one of the main predictors of species vulnerability to extinction (Harcourt et al. 2002; Munday 2004; Colles et al. 2009; Sagot & Chaverri 2015). *T. tricolor* is already known to be a highly specialized bat that only uses tubular leaves for roosting. Our current results show that the available plant species may also highly limit this species distribution, within that already narrow niche. Suppose biomechanical constraints impose potential costs to using leaves without a prominent apex. In that case, we could predict that either the bats suffer accidents constantly or learn to more adequately approach and enter less suitable leaves at sites where the preferred roost is rare or unavailable. Regardless of the answer, our results show that biomechanics should be incorporated into resource selection studies, especially when complex manoeuvres are needed to acquire those resources.

## ACKNOWLEDGEMENTS

We would like to thank Silvia Chaves-Ramírez and Mariela Sánchez-Chavarría for assistance during field data collection. Julio Bustamante and Lilliana Rubí Jimenez provided valuable guidance during research permit application. We also thank Ronald Villalobos for logistics support and the Centro Biológico Hacienda Barú for their continuous support of our research.

## Data availability

The data and code supporting this article will be available from the GitHub repository.

## Author contributions

Conceptualization: G.C, M.A.S., J.A.; Methodology: G.C, M.A.S., J.A.; Investigation: G.C., J.P.B., T.U.E., M.P.A., A.L.V., J.A.; Formal analysis: M.A.S., G.C., J.A.; Resources: G.C., J.A.; Data curation: M.A.S., G.C., J.A.; Writing - original draft: G.C., M.A.S.; Writing - review & editing: G.C, M.A.S., J.A., J.P.B., T.U.E., M.P.A., A.L.V.; Visualization: M.A.S.; Supervision: G.C.; Funding acquisition: G.C., J.A.

